# Collective intercellular communication through ultra-fast hydrodynamic trigger waves

**DOI:** 10.1101/428573

**Authors:** Arnold J. T. M. Mathijssen, Joshua Culver, M. Saad Bhamla, Manu Prakash

**Author notes:** Please address correspondence to: M.S.B or M.P.

## Abstract

The biophysical relationships between sensors and actuators [1–5] have been fundamental to the development of complex life forms; Abundant flows are generated and persist in aquatic environments by swimming organisms [6–13], while responding promptly to external stimuli is key to survival [14–19]. Here, akin to a chain reaction [20–22], we present the discovery of hydrodynamic trigger waves in cellular communities of the protist *Spirostomum ambiguum*, propagating hundreds of times faster than the swimming speed. Coiling its cytoskeleton, *Spirostomum* can contract its long body by 50% within milliseconds [23], with accelerations reaching 14g-forces. Surprisingly, a single cellular contraction (transmitter) is shown to generate long-ranged vortex flows at intermediate Reynolds numbers, which can trigger neighbouring cells, in turn. To measure the sensitivity to hydrodynamic signals (receiver), we further present a high-throughput suction-flow device to probe mechanosensitive ion channel gating [24] by back-calculating the microscopic forces on the cell membrane. These ultra-fast hydrodynamic trigger waves are analysed and modelled quantitatively in a universal framework of antenna and percolation theory [25, 26]. A phase transition is revealed, requiring a critical colony density to sustain collective communication. Our results suggest that this signalling could help organise cohabiting communities over large distances, influencing long-term behaviour through gene expression, comparable to quorum sensing [16]. More immediately, as contractions release toxins [27], synchronised discharges could also facilitate the repulsion of large predators, or conversely immobilise large prey. We postulate that beyond protists numerous other freshwater and marine organisms could coordinate with variations of hydrodynamic trigger waves.

## INTRODUCTION

Trigger waves solve two fundamental problems caused by diffusion in biological signalling; they do not slow down nor lose amplitude, allowing to reliably communicate over large distances [20–22]. These feed-forward wave fronts are often chemical in nature, such as neuronal spikes at ~ 100m/s or Ca^2+^ waves at ~ 30*µ*m/s [22], but require an excitable medium (cellular membrane or cytoplasm) or physical contact (neurons). On the one hand, fluid media do not feature such constraints, facilitating endocrine transport (adrenaline release) at ~ 30mm/s [22] or cytoplasmic streaming at ~ 100*µ*m/s [28], but generally need minutes for full recirculation, decay with distance or size, and fundamentally lack midway signal reinforcement. On the other hand, hydrodynamic interactions can mediate collective motion in ‘living fluids’ [8–11], but much less is known how this links with biological sensing and decision making [5, 14–19].

## ULTRA-FAST CONTRACTION KINEMATICS

*Spirostomum ambiguum* is one of the largest unicellular protozoans, with body length *L*_0_ = (1.1 ± 0.2)mm [Fig. 1A,B]. By coordinating its cilia in metachronal waves [Movie S1] it swims at low Reynolds numbers, 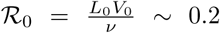, with speed *V*_0_ in fresh water of viscosity *ν* = 1mm^2^/s [Movie S2]. However, as a defence against predators, the cell can rapidly contract to a fraction *f_c_* = 0.4 ± 0.03 of its length within *τ_c_* = (4.64 ± 0.15)ms [Fig. 1A,C & Movie S3]. Then, as the Reynolds number surges to 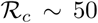, it releases toxins [27]. After contraction the cell slowly relaxes, in *τ_r_* ~ 1s. A pictorial summary of the underlying biochemical mechanisms is given in Fig. 1D and SI §2A.

**FIG. 1.**
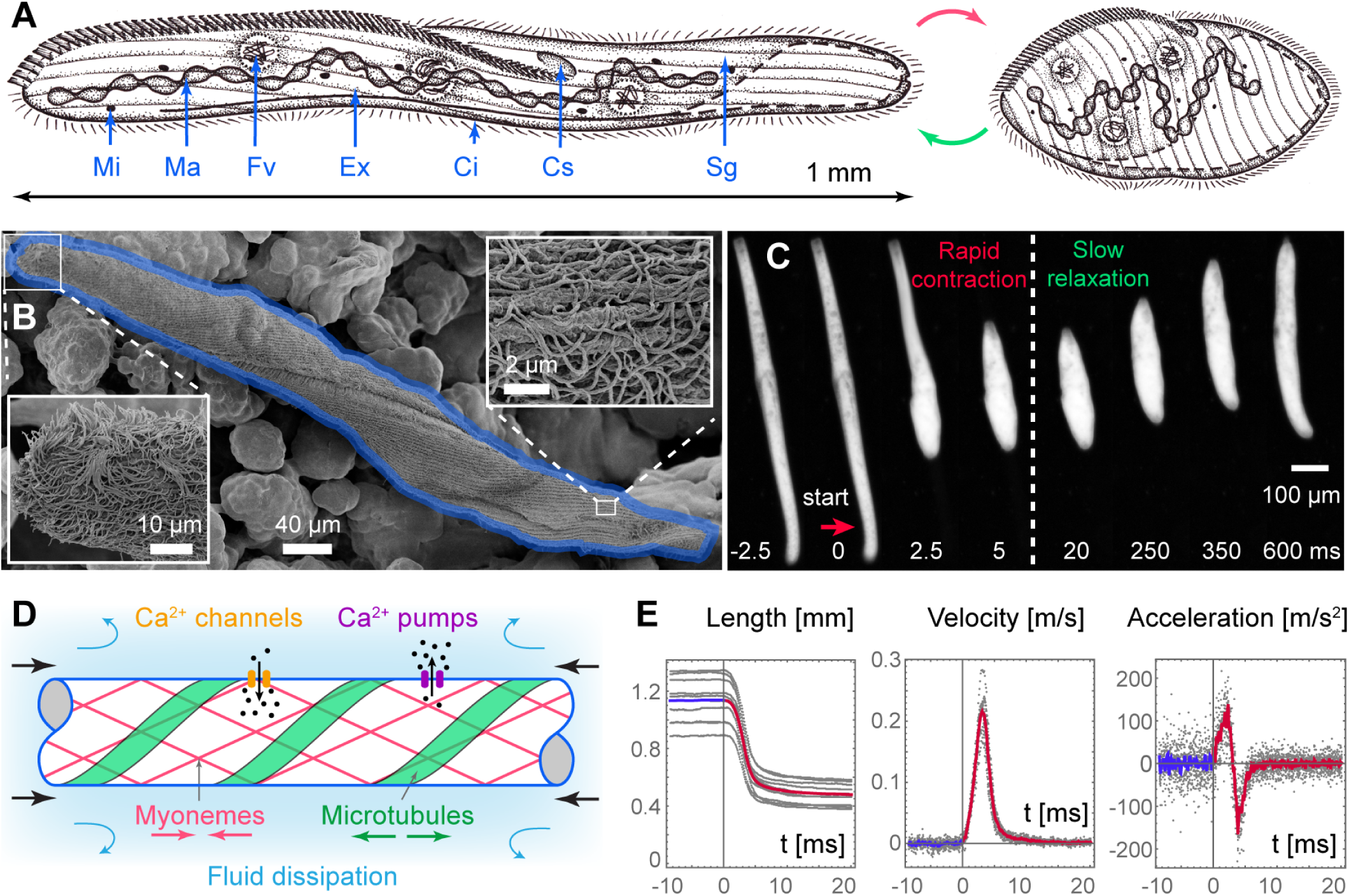
Spirostomum’s contraction dynamics. **A**. Anatomy diagram: Micronuclei (Mi), Macronuclear nodules (Ma), Food vacuoles (Fv), Extrusomes containing toxins (Ex), Cilia (Ci), Cytostome (Cs), Somatic grooves (Sg). **B**. SEM micrograph. Insets highlight cilia and somatic grooves. **C**. Collage of a spontaneous contraction and recovery. **D**. Biochemistry diagram of antagonistic contraction and relaxation mechanisms [SI §2A]. **E**. Kinematics of organism length *L*, velocity 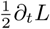 and acceleration 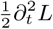. Shown are average values (red) for ten different organisms (gray) triggered by electric stimulation (|***E***| = 1.5kV/m), with contraction onsets aligned at *t* = 0.

To quantify these dynamics systematically, we developed an electrical stimulation apparatus [23] with microsecond precision, synchronised with high-speed (10,000 fps) microscopy [SI §2]. Figure 1E shows the cell length, velocity and acceleration, and from statistics of thousands of cells we construct the complete phase diagram of contractile behaviour [Fig. S1]. The velocity peaks at *V*_max_ = (0.22 ± 0.01)m/s, and the acceleration at *A*_max_ = (139±28)m/s^2^ ~ 14 g-forces. Unlike other rapid cellular movements like nematocysts firing [29], *Spirostomum* is capable of this actuation repeatably - over and over again - making it a fascinating model organism to study ultra-fast motion in biology [4].

## FLOW GENERATION

The contraction generates long-ranged flows around the organism, which we analyse in a liquid film experiment [Fig. 2A]. The measured flow structure is contractile along the cell’s major axis [Figs. 2C,D & Movie S4], resembling the ‘puller-type stresslet’ often produced by micro-organisms [7, 10, 13]. A dramatic difference, however, is that vortices emerge that expand into the medium over time. This vortex generation is a signature of inertia, also apparent as a delay in fluid motion [Fig. 2E], since the boundary layer width grows as 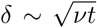 [15]. Indeed, conducting control experiments by suspending *Spirostomum* in high-viscosity medium shows no such delay, and no vortices, in agreement with the Stokesian stresslet (*ν* = 50mm^2^*/*s and *τ_c_* = 15ms, so 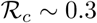).

**FIG. 2.**
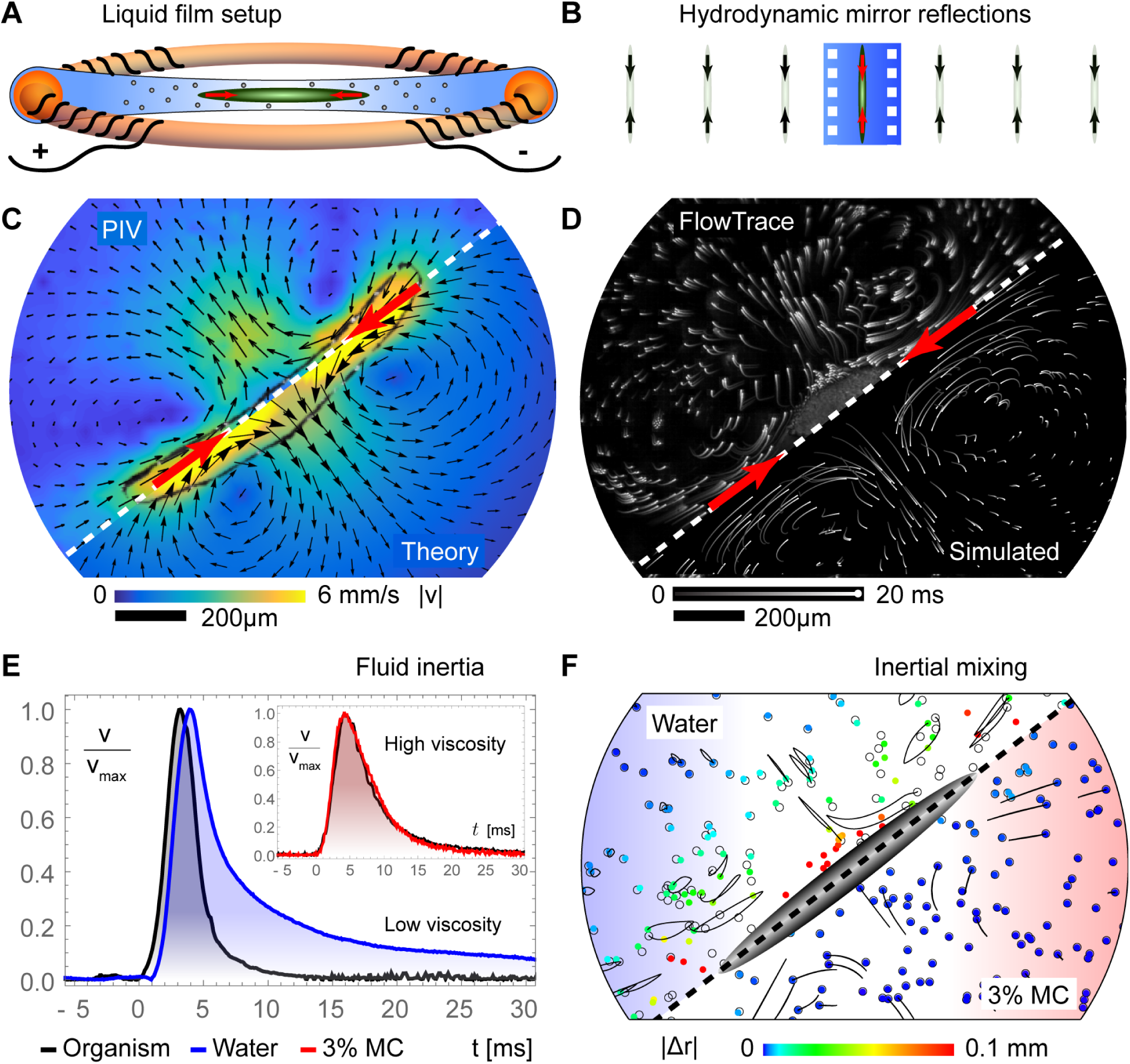
Flow generation during contractions. **A**. Diagram of experimental setup. A liquid film is suspended in a ring, with two electrodes. **B**. Diagram of hydrodynamic image systems reflected by water-air interfaces. **C**. Comparison of PIV measurements and LNS theory (Eq. 1), averaged over 40ms after the onset of contraction. **D**. Comparison of experimental and simulated tracer dynamics, 20ms after contraction onset. **E**. Offset in organism boundary (black) and flow velocities, at low viscosities (water, *ν* = 1mm^2^*/*s, blue) and high viscosities (3% methyl cellulose, *ν* = 50mm^2^*/*s, red). **F**. Displacement of tracer particles after contraction and relaxation, simulated for low (blue) and high (red) viscosities.

Therefore, to model the contraction flow with inertia, we solve the linearised Navier-Stokes (LNS) equations [12] combined with the method of images [13] to account for the liquid-air interfaces [Fig. 2B & SI §3]. We extend the stresslet to a set of equal and opposite point forces, ***f****_k_*(*t*′), distributed along the cell, giving the flow

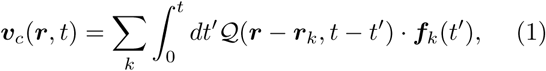

where 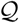 is the Green’s function tensor with inertial memory kernel, given explicitly in SI §3B. Importantly, both the spatial and temporal elements of this model agree with the measurements, especially how the vortices expand [Fig. 2C & Movie S5]. We also recover the particle dynamics when integrating the flow over time [Fig. 2D].

Moreover, the effect of inertial mixing is highlighted by simulating the organism’s contraction followed by slow relaxation [Fig. 2F, SI §3C,]. Traditionally, the ‘scallop theorem’ forbids viscous mixing by any reciprocal motion [6]. However, using inertia *Spirostomum* can displace particles up to 10% of its body length. This effect could further facilitate the dispersal of toxins and enhance food transport [3], significantly out-competing thermal diffusion with Péclet numbers 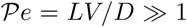 even for the smallest molecules (where *D* is the diffusion constant).

## RHEOSENSING

To detect predators, organisms must sense surrounding flows. As the liquid exerts hydrodynamic stresses [14], the cell membrane stretches and hence mechanosensitive ion channels open [18, 24]. To probe *Spirostomum’s* rheosensitivity, we developed a high-throughput (~ 10cells/s) microfluidic assay [Fig. 3A]. In this radially symmetric device we introduce organisms on the outer edge and apply a suction flow drawing liquid to the central outlet. The resulting flow stretches the organisms, with strain rate 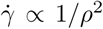 at position *ρ* [SI §4A]. This geometry was inspired by ‘spaghettification’ [30], a stretching induced by tidal forces near a black hole, rendered here microscopically [SI §4B]. Therefore, as cells are pulled towards the central region, the strain increases until they reach a threshold 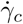 and contract [Movie S6]. Imaged from below, we observe that all cells are triggered around a ring of radius *ρ_c_*, the ‘event horizon’ [Fig. 3B]. Ergo, we obtain the critical strain rate, 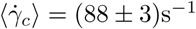.

**FIG. 3.**
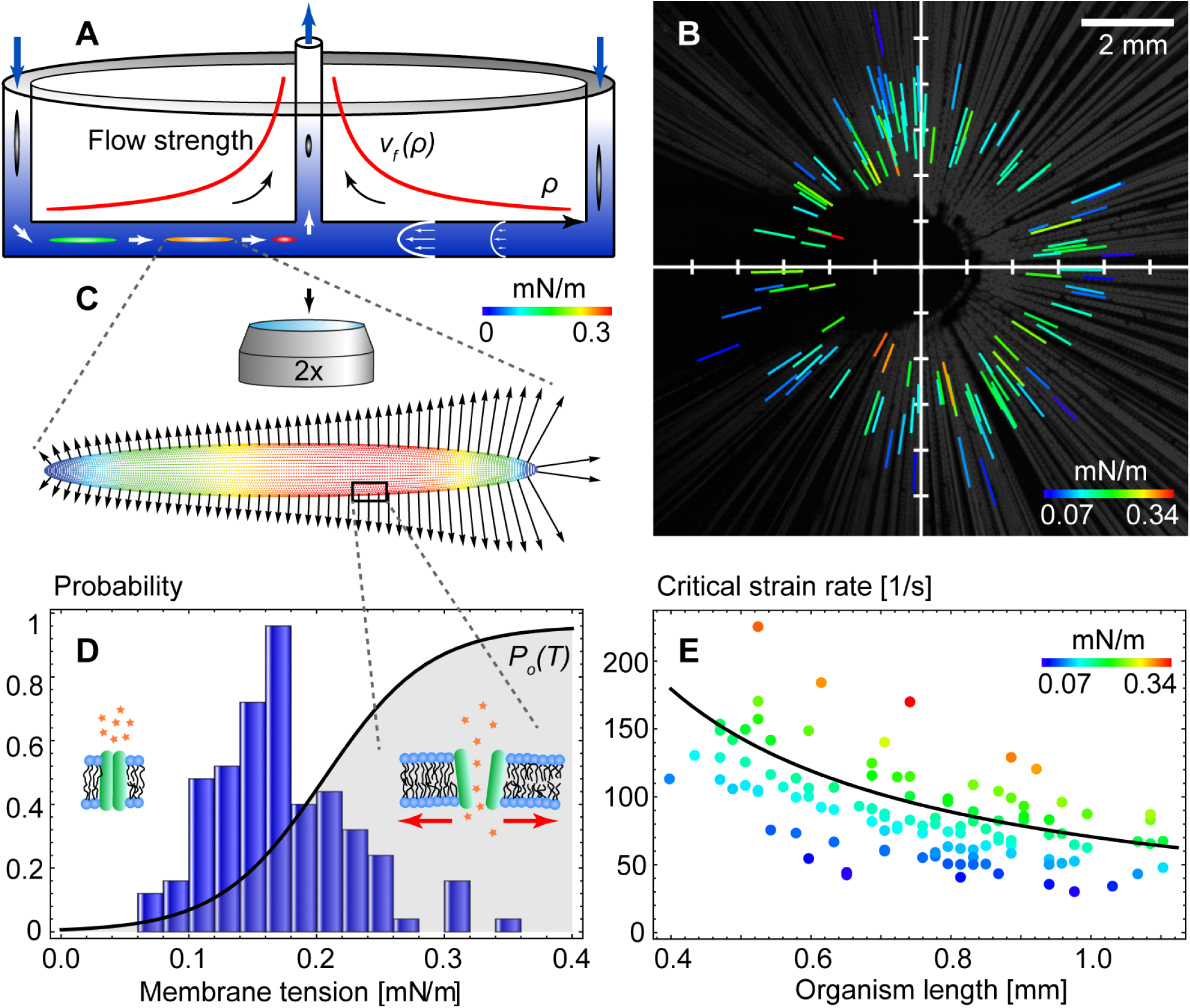
Rheosensing experiments. **A**. Diagram of microfluidic set-up. Inset: Flow strength vs. radial distance. **B**. Gray lines are organism trajectories as they move towards the central outlet. Coloured lines show their maximum membrane tension and position at the onset of contraction, *N* = 115. **C**. Hydrodynamic forces (arrows) and tension (colours) on the cell membrane, calculated with the MRS. **D**. Measured distribution of critical membrane tension, pdf(*T_c_*) (blue), and theory for the gating probability of a mechanosensitive ion channel (black). **E**. Critical strain rate and tension versus organism length, 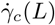, as measured (colours) and theoretical prediction (black line).

Crucially, using this macroscopic flow we can back-calculate microscopic forces that act on mechanosensitive ion channels. Employing the Method of Regularised Stokeslets (MRS) [31], for each organism, we compute the force distribution on the membrane at the moment before contraction [Fig. 3C and SI §4C]. Hence, we find the critical membrane tension required for the ion channels to open, 〈*T_c_*〉 = (0.17 ± 0.02)mN/m [Fig. 3D]. This assay can be applied straightforwardly to other organisms with different sizes or more complex shapes.

**FIG. 4.**
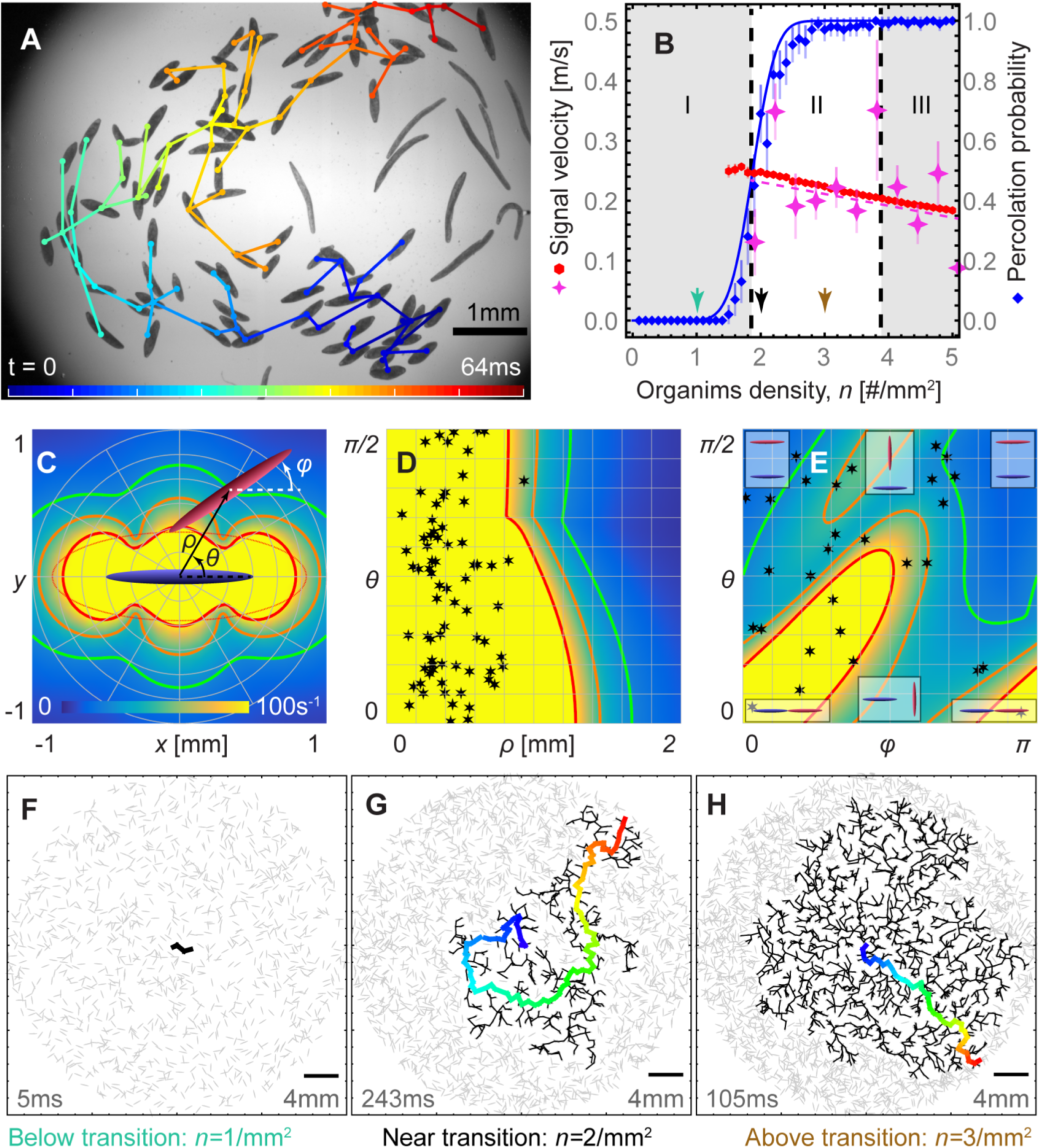
Collective hydrodynamic communication. **A**. Trigger wave propagating though a colony. The graph indicates which organism triggered which. **B**. Phase diagram. Percolation probability, *P*(*n*), in simulations (blue diamonds) and theoretical prediction (blue line). Superimposed are wave velocities, *v_w_*(*n*), in simulations (red) and experiments (pink stars). **C-E**. Pairwise interactions between transmitter *A* (blue) and receiver *B* (purple). **C**. Generated strain rate (Eq. 1), and the sensing threshold 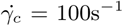 (solid red). This threshold is approximated by an ellipse (dashed red). **D**. Strain rate on the body of *B* for relative positions (*ρ, θ*), maximising over all orientations *ϕ*. Stars show corresponding experimental observations. **E**. Same, for relative orientations (*ϕ, θ*), with constant *ρ* = 1mm. **F-H**. Simulations of trigger waves in colonies below, near, and above *n_c_*. Gray ellipses are non-triggered organisms. Black lines show the triggered connectivity graph. The first percolating path to the colony edge is shown with colours scaled to the percolation time (bottom left corners).

These measurements are consolidated with established theoretical results for ion channel gating [24]. Using the ‘two-state’ model, the probability for a channel to open, *P_o_*(*T*), is estimated as a function of applied tension [SI §4D]. Indeed, this probability transitions at a threshold tension 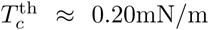 that agrees well with the experimental values. This result underlines the significance of the spatial distribution of hydrodynamic stresses, and allows us to predict the stimulation threshold for other organisms or geometries.

In particular, cell size is expected to contribute a major role in rheosensitivity. When mapping the observed critical strain rate 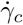 against length, we find that larger cells require smaller stimuli to contract [Fig. 3E], as also predicted theoretically [SI §4E]. This suggests that longer organisms are better sensors, likely being an evolutionary advantage in the endeavour to discern predators, which could elucidate why *Spirostomum* is one of the longest known ciliates and takes its unusual cigar shape.

## HYDRODYNAMIC COMMUNICATION

To ascertain the physiological role of hydrodynamic trigger waves, we observe *Spirostomum* in undisturbed growth chambers over long periods of time. Surprisingly, *Spirostomum* tends to self-assemble into clusters, as reproduced by mixing cultures and then recording accumulation with time-lapse imaging [SI §5A & Movie S7]. Once close together, the organisms can exhibit a remarkable collective behaviour: when a first cell spontaneously contracts, it generates a flow that can trigger neighbours, and hence initiate a cascade of contractions that propagates through the colony [Fig. 4A, SI §5B & Movie S8]. These hydrodynamic trigger waves travel at remarkable speeds, *v_w_* ~ 0.25m/s [Fig. 4B], not inappreciable compared to human neurotransmission [22].

We examine the communication potential by first considering pairwise interactions [Fig. 4C & SI §5C]. When organism *A* (blue) transmits a signal (Eq. 1), organism *B* (purple) responds if it crosses the red line, 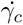. This model agrees with pairwise experiments for different relative configurations (*ρ, θ, φ*) [Fig. 4D,E], showing that not only the distance but also orientations matter. Specifically, ‘one-way streets’ can form where a signal can only travel from *A* to *B*, e.g. 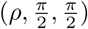 versus 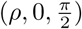, as the flow is inherently directional. We quantify this with antenna theory [SI §5D], originally designed for electromagnetic radiators, showing the hydrodynamic signals are ~ 2x stronger along the body axis (directive gain 2.79 dB).

Using these building blocks, we consider multi-organism interactions by simulating large colonies of different cell densities *n* [SI §5E]. The central cell is triggered at time *t* = 0, and cascades computed subsequently [Fig. 4F-H]. At low densities (*I*, 1 mm^−2^), neighbours cannot be reached sufficiently so the signal rapidly dies out. At high densities (*II*, 3 mm^−2^), signals propagate radially outwards and abundantly reach the colony edge. In between, near the critical density 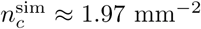, the signal does not travel radially but in fractals, much like lightning discharge patterns. Here the probability for a signal to reach the colony edge, *P*(*n*), transitions rapidly from zero to one [Fig. 4B]. Importantly, a small change in density therefore has a large impact on the communication potential near criticality [Movie S9].

This phase transition is understood in terms of percolation theory [25]. As detailed in SI §5F, the percolation threshold is estimated analytically as 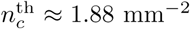. Hence, this theory offers a fair agreement with our simulations, and also with our experiments where signals are only sustained above 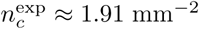 [Fig. 4B]. Moreover, the theory predicts a lower percolation threshold for higher cell aspect ratios, again highlighting *Spirostomum*’s slender body.

Yet, at the highest densities, (*III*, 5 mm^−2^), the wave velocity *v_w_*(*n*) reduces. The signal is delayed because every organism (per unit area) requires time to perceive and respond, as demonstrated in Movie S10. Too high densities are also disadvantageous in terms of energy expenditure and sharing food reserves. Consequently, organisms are driven to criticality, to *n_c_*, when optimising for resources and conductivity.

## DISCUSSION

To conclude, hydrodynamic communication could be advantageous to organise cellular communities, either for the defence against large predators [27] by synchronised toxin discharge, or conversely to immobilise large prey [32], or for collective nutrient mixing. Beyond immediate benefits the signalling could also regulate long-term behaviour, influencing gene expression. However, the individual judgement to contract must be gauged carefully, for energy efficiency but also because the extreme forces in repeated contractions can potentially damage cells. Certain aspects of this decision-making process may therefore be compared with quorum sensing [16], considering that signals are sustained only above a critical organism density.

The presented percolation theory can be extended to more natural 3D organism colonies using ellipsoids of revolution. However, on a more fundamental level, we expect hydrodynamic trigger waves to be part of the *directed percolation* universality class [26]. Directional symmetry breaking, like following ‘one-way streets’ in a maze, is also observed here. Pursuing this could be an exciting new avenue for future research in biological critical phenomena.

Finally, similar rapid contraction mechanisms are observed in other model ciliates including *Vorticella* [1] and *Stentor* [5]. Beyond protists, innumerable organisms both generate and sense flows, from bacteria to fish [2–7, 12–19], which opens a scope for new questions how broadly hydrodynamic trigger waves could contribute to collective behaviour [8–11] and, more generally, to science of active matter and adaptive materials.

## SUPPLEMENTARY MATERIALS

- Supplementary Information (.pdf)
- Supplementary Movies S1 – S10 (.mov)

## ACKNOWLEDGEMENTS

The authors would like to thank Scott Coyle and Deepak Krishnamurthy for helpful discussions. A.M. acknowledges funding from the Human Frontier Science Program (Fellowship LT001670/2017). This work was supported by NSF CCC grant (DBI-1548297) (M.P.), US Army Research Office grant (W911NF-15–1–0358) (M.P.), CZI BioHub Investigator Program (M.P.), the Howard Hughes Medical Institute (M.P.), and NSF grant (award number 181733) (M.S.B). We thank M. Gruber for the scientific illustration of *Spirostomum ambiguum* shown in Figure 1a.

## AUTHOR CONTRIBUTIONS

A.M., M.S.B., and M.P. designed the research, A.M., M.S.B. and J.C. performed the experiments, A.M. analysed the data, A.M. carried out the simulations and theory, A.M., M.S.B., J.C. and M.P. wrote the manuscript.

## COMPETING INTERESTS

The authors declare no competing interests.

## Supplementary Information

### I. ORGANISM CULTURING

*Spirostomum ambiguum* (Sciento catalog #P346) was cultured in Hummer medium [1] of spring water infused with Timothy hay (Carolina catalog #132385) and boiled wheat grains (Carolina catalog #132425). Cultures were incubated at 18^◦^C in dark conditions, and reinoculated fortnightly. Organisms used in experiments were extracted in the exponential growth phase, rinsed and transferred to fresh medium.

### II. ULTRA-FAST CONTRACTION KINEMATICS

#### A. Biochemical mechanism

Although the exact molecular mechanisms [2–8] of contraction and relaxation is not fully understood, the key elements may be summarized as follows [Fig. 1D of the main text]; At rest, the cell maintains a low internal Ca^2+^ concentration using ion pumps (purple). When triggered, electro-or mechanosensitive ion channels (orange) open to raise the Ca^2+^ concentration rapidly. These ions bind to engage ‘myoneme’ filaments (red) that generate tensile forces. The rapid acceleration is halted by the cytoskeleton (green) building up elastic energy and fluid dissipation (blue). To recover, the ion pumps reduce the Ca^2+^ concentration to homeostasis, so the myonemes relax and the elastic microtubules restore elongation.

#### B. Experimental details

The dynamics of *Spirostomum* contractions are measured with high-speed (Phantom V1210 camera, 10,000 fps) microscopy (Nikon Te2000 scope, 1.5X objective), as shown in [Fig. S1A]. To collect a large amount of statistics, organisms are stimulated electrically with an electrophysiological DC pulse generator (Grass instruments, S88). The organisms are introduced in a rectangular microfluid chip (*L_c_* =2.5cm, *W_c_* = 1cm, *H_c_* = 200*µ*m) of glass slides with copper side walls that act as electrodes. For a given DC voltage *V*, the applied electric field strength is |***E***| = *V/W_e_* and the pulse duration is *τ_E_*. The pulse and camera trigger are synchronized with a micro-controller (Arduino Uno) and monitored separately with an oscilloscope (Instek GDS-1152A-U).

**FIG. S1.**
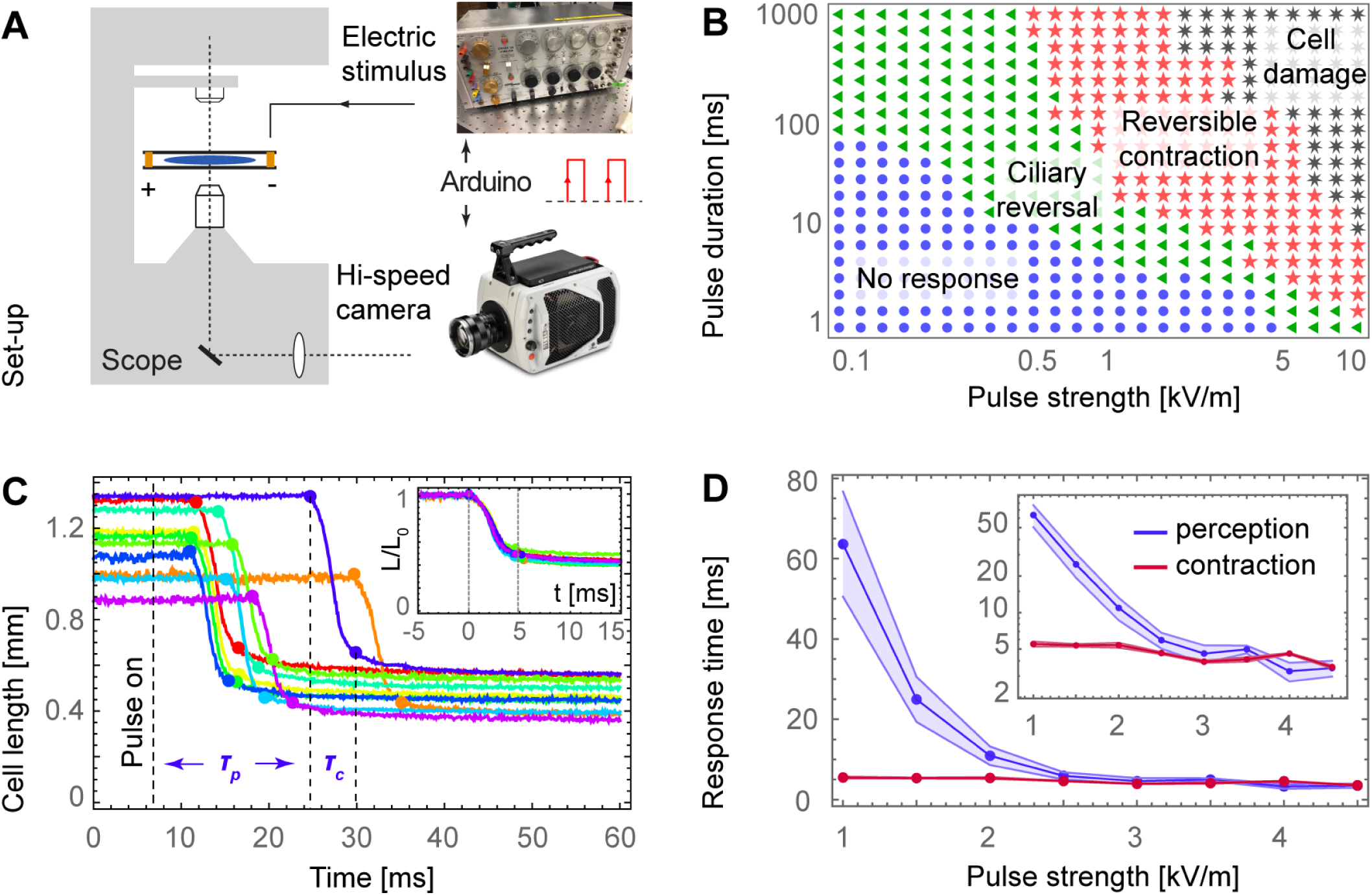
Measurement of electrically stimulated *Spirostomum* contraction dynamics. A. Diagram of experimental set-up. B. Phase diagram of organism response to electric signals. Shown is median behaviour of *N>* 20 organisms for each data point. C. Typical kinematics of organism length after stimulation at |***E***| =1.5kV/m and *τ_E_* = 100ms. Inset: Collapse of kinematics, when normalising the cell length and lining up the onsets of contraction at *t* = 0. D. Perception and contraction time against stimulus intensity, with *τ_E_* = 100ms. Shaded regions are error bars of 95% CL. Inset: same on log scale.

We first map the organism response to different stimuli by varying the pulse duration *τ_E_* and electric field strength |***E***| [Fig. S1B]. Note, for each data point the channel is rinsed and *N >* 20 new organisms are introduced. For mild stimuli the organisms temporarily swim backwards through ciliary reversal, a well established behaviour [9, 10]. Intermediate stimuli induce reversible contractions, as observed in natural conditions with non-electric stimuli, but strong pulses lead to cell damage. Therefore, we perform all further electric experiments in the intermediate regime, with *τ_E_* = 100ms unless otherwise stated.

Typical dynamics for |***E***| =1.5kV/m are shown for 10 organisms in [Fig. S1C]. The cell lengths *L*(*t*) are found by analyzing the images with MATLAB built-in edge detection and ellipse fitting. Subsequently, the velocity *V* (*t*)= *∂_t_L* and acceleration 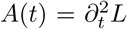 are computed, as shown in Fig. 1 of the main text.

Repeating this experiment for different voltages, we extract the perception time, the period between the pulse and contraction onset *τ_p_*, and the contraction duration *τ_c_* [Fig. S1D]. The perception time is large for weak pulses, *τ_p_* = (64 ± 13)ms at |***E***| = 1kV/m, but short otherwise, *τ_p_* = (3.5 ± 0.5)ms at |***E***| *>* 2.5kV/m. This reflects the strength of ion fluxes [SI §2A], in agreement with [11]. The contraction time is approximately constant for all stimuli, *τ_c_* = (4.64 ± 0.15)ms, as measured for *N* = 199 organisms, reflecting the time required for the myonemes to engage. As a result, the contraction dynamics collapse onto a single curve [Fig. S1C, inset].

#### C. Kinematics model

Combining the information from these experiments, we construct a simple model for the contraction kinematics. The organism length can be described as

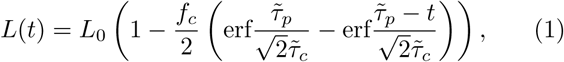

where the initial body length is *L*_0_ ≈ 1mm, the fraction of length contraction is *f_c_* ≈ 0.5 so the final body length is *L*_0_ *f_c_*, the perception time is *τ_p_* ≈ 3.5ms and the contraction duration is *τ_c_* ≈ 5ms, with 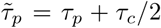 and 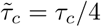. Subsequently, the organism velocity is

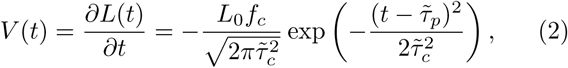

which is a Gaussian peaked halfway the contraction. We will use this model in the next sections.

We define the Reynolds number during contraction as

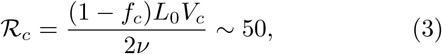

using typical length scale *L*_0_ ≈ 1mm, *V_c_* = *L*_0_*/τ_c_* ≈ 0.2m/s, and contraction fraction *f_c_* ≈ 0.5, where the factor 2 accounts for the fact that *V_c_* constitutes both the head and tail end velocities moving towards one another.

### III. FLOW GENERATION

#### A. Experimental details

The flows generated during a contraction is imaged with high-speed (Phantom V1210 camera, 10,000 fps) microscopy (Nikon Te2000 scope, 10X objective), under dark-field illumination. The experimental geometry is a liquid film, inspired by bacterial mixing experiments [12, 13]. The film is suspended in an acrylic ring (insulator) of diameter *D* = 1cm and height *H* = 500*µ*m, with thin copper wire electrodes wrapped on either side [Fig. 2A]. Organisms are taken from cultures in the exponential growth phase, resuspended in fresh medium, and polystyrene tracer particles (Polybead catalog #17141–5, diameter 6.0*µ*m) were added at concentration *ϕ* ~ 10^−2^% solids (w/v). These beads feature a coating of carboxyl groups to prevent adhesion to the organisms. A volume *V* = *πD*^2^*H/*4 ~ 393*µ*L of this suspension, including one organism, is pipetted into the ring. The organism is free to swim, and triggered (|***E***| = 1.5kV/m, *τ_E_* = 100ms) when located at the centre of the ring.

The contraction flow is shown in Movie S4, where the temporal dynamics are highlighted with FlowTrace [14]. These currents are is analysed with PIVlab [15] for cross-correlation and TrackMate [16] for direct particle tracking. The PIV results are compared with our theory [see next section] in Fig. 2C. The particle trajectories are used to calculate the flow speed, by averaging over *N* = 171 tracers. In Fig. 2E this is compared to the organism speed, found by taking the first derivative of the organism length. The fluid inertia causes a delay between the liquid and its boundary condition, being the moving organism surface.

As a control for inertia, we repeat our experiments in a high-viscosity (*ν* = 50mm^2^*/*s) solution of 3% Methyl Cellulose (Sigma Aldrich catalog #M7140) dissolved in culture medium. Because of its polymeric nature this agent is mostly inert to the cell biochemistry. The organisms contract slower in this solution because of the higher viscosity, *τ_c_* = 15ms, so the Reynolds number is reduced further down to 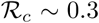. Hence, the flows are Stokesian, without inertial delay and without vortexes spreading into the liquid.

#### B. Contraction flow model

We consider an organism located at the origin, at position (*x, y, z*) = (0, 0, 0) in Cartesian coordinates. The long axis of the cell is oriented in the 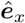 direction.

##### 1. Stokes flow

First, in the absence of inertia, the hydrodynamics of an incompressible flow at Reynolds number 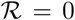 are described by the Stokes equations,

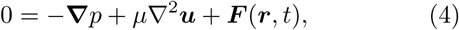

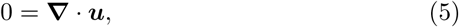

where ***u***(***r****,t*) is the flow velocity at position ***r*** and time *t*, *p*(***r****,t*) is the pressure, *µ* is the dynamic viscosity and ***F*** (***r****,t*) is a force acting on the liquid. We first consider the flow in the absence of surfaces, with boundary conditions ***u*** = 0 as |***r***|→∞. The fundamental solution to these equations, the flow due to a point force ***F****^S^*(***r***, *t*) = *δ*^3^(***r*** − ***r***′)***f***(*t*), is called the Stokeslet

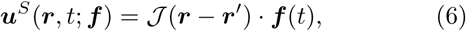

where the Oseen tensor 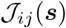 has Cartesian components

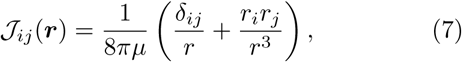

with *i, j* ∈{*x, y, z*} and *r* = |***r***|. This solution is shown in Fig. S2A. Because the point force moves during the contraction, the position ***r***′ = ***r***′(*t*) depends on time. However, as the Stokes equations only depend on time through the force ***F*** (***r****,t*) and the boundary conditions, the flow adapts instantaneously.

##### 2. Superposition

Because the equations are linear, the flow due to a number *N_k_* of point forces, ***F****_k_* = *δ*^3^((***r*** − ***r****_k_*)***f****_k_*(*t*), can be written as the superposition

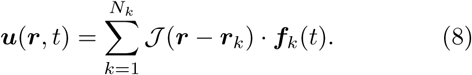

Specifically, similar to electrostatics, one can perform a multipole expansion of the Stokeslet flow [17]. The ‘Stokes dipole’ or ‘Stokes stresslet’ is the combination of two equal and opposite forces, ‘Stokes dipole’ or ‘Stokes stresslet’ is the combination of two equal and opposite forces,

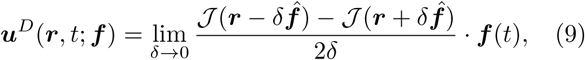

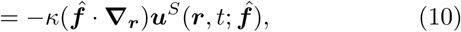

where *κ* =2δ|***f***| is the dipole strength and the dimensionless direction 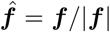. This flow is already a first approximation of the *Spirostomum* contraction.

**FIG. S2.**
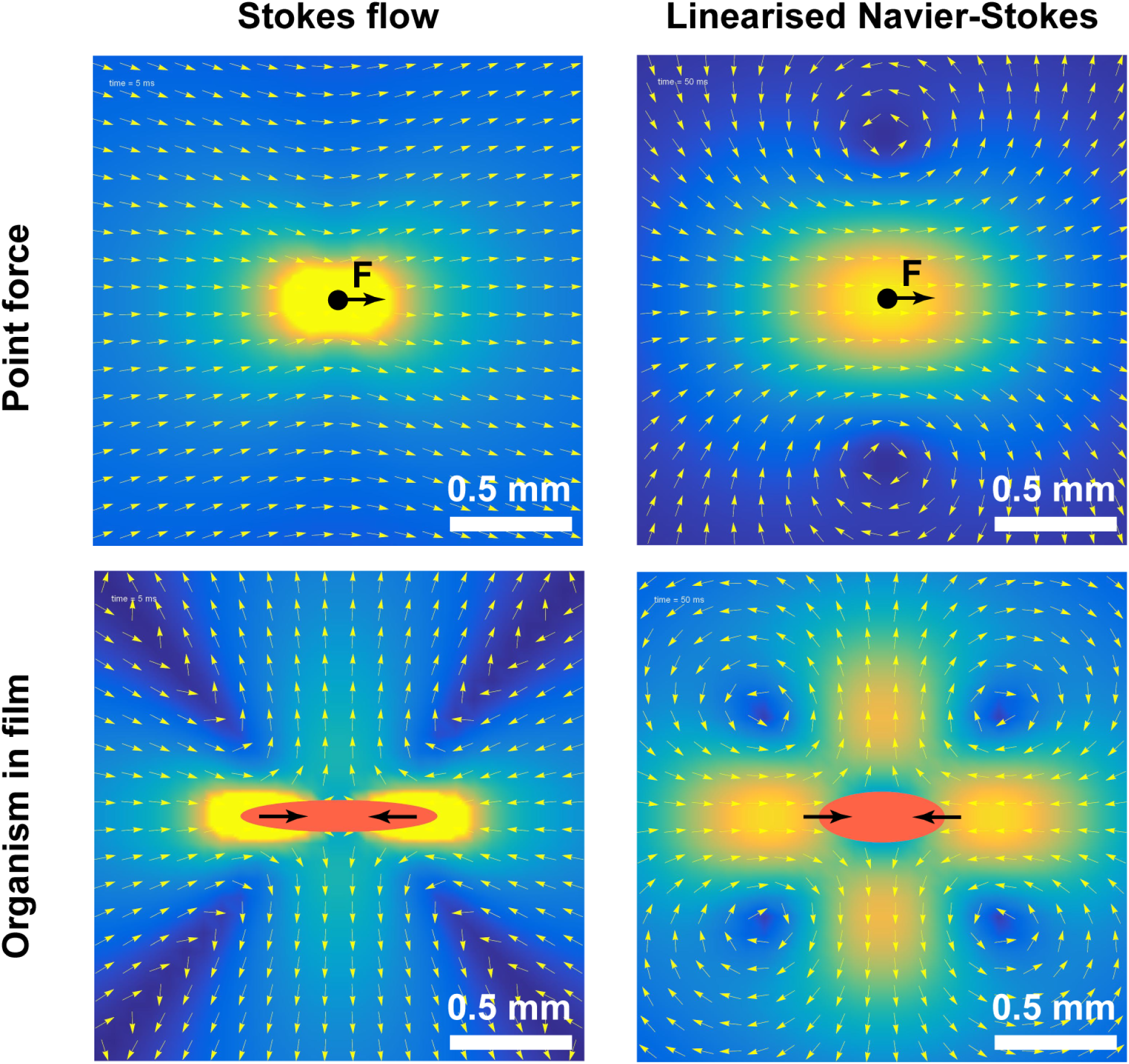
Comparison of Stokes flow 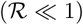 and Linearised Navier-Stokes flow 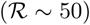. A. Point force solution; Stokeslet (6). B. LNS tensor solution (14). C-D. Contraction flow in a liquid film (20); for high (C; 1000x water) and low (D; water) viscosities, with *m* = ±100 image reflections and *k* = ±5 grid points on the elongated body. Cf. Movie S5. The Stokes solution has open stream lines but the LNS solution features vortices that spread out over time. Colours show flow magnitude and arrows are stream lines.

##### 3. Liquid film confinement

To account for the confinement of the experimental liquid film geometry, we must satisfy the no-shear-stress boundary condition at the liquid-air interfaces, *z* = ±*H/*2 for film height *H* = 500*µ*m. Therefore, we employ the method of images [18], again borrowed from electrostatics. These images act like ‘mirror reflections’ of the applied forces. Now imagine walking in a hall of mirrors. Because we have two liquid-air interfaces there will be a reflection of the reflection, a reflection of the reflection of the reflection, etc. Thus, the resulting flow can be expressed as an infinite series. For forces located in the middle of the film, at *z* = 0, the images are located at *z* = ±*H*, ±2*H*, ±3*H*, …. Hence, it can be shown that the solution for a dipole aligned along 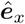 and centered in the film is

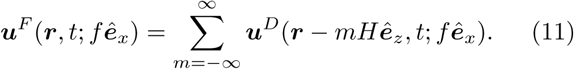

This solution converges, as the dipole flow decays with distance as 1*/r*^2^. Hence, the series can be truncated to ±*M* terms for distances *r* ≪ *MH*. We will simulate flows up to *r* ~ 5mm, so we truncate at *m* = ±100.

There are two points of caution that must be considered before continuing. First, an infinite series of Stokeslet forces does not converge unconditionally because it decays as 1*/r*, so more care must be taken. However, we are only concerned with organism contractions without external forcing. Second, for an experimental geometry between glass sides, instead of a liquid film, the image systems are more complex [19]. In this case, the boundaries can induce recirculation even at zero Reynolds number [20]. This recirculation could be confused with the emergence of inertial vortices. But this is not the case for our dipolar flows in a liquid film [18], which feature vortices only at 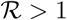 but no vortices and open stream lines in high-viscosity medium.

##### 4. Inertial flow

To account for effects of inertia, we consider the linearized Navier-Stokes (LNS) equations [21–23].

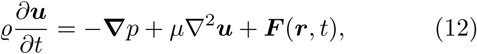

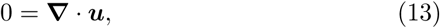

where 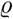 is the fluid density, so the kinematic viscosity is 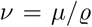. The ‘oscillatory Reynolds number’ associated with these equations is 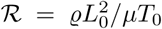, where *L*_0_ and *T*_0_ are typical length and time scales. The fundamental solution to these equations can also be expressed in terms of a hydrodynamic tensor,

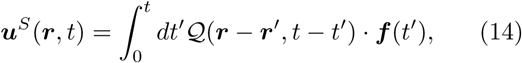

where the integration over the memory kernel arises because this inertial flow now requires a period 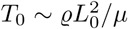 to adapt to changes in forcing. The explicit expression of the hydrodynamic tensor is

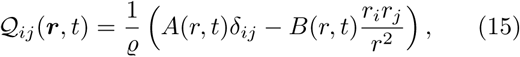

in terms of the scalar functions

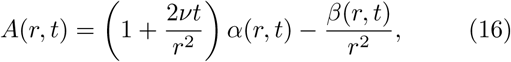

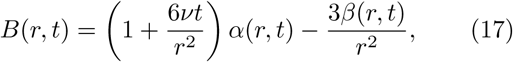

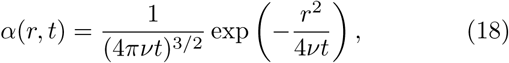

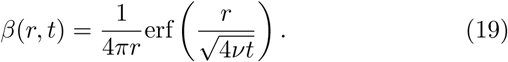

Indeed, in the high-viscosity limit, when the tensor 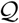 varies much more slowly in time than the forcing ***f****_k_*, we recover 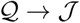, the Oseen tensor (7).

##### 5. Combined flow

Combining this LNS solution (14–19) with the concepts of superposition (8) and the method of images (11), we model the *Spirostomum* contraction flow as

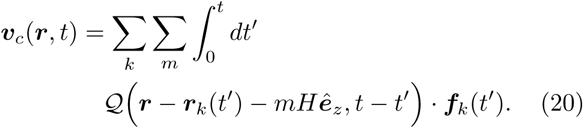

The equal and opposite forces that the organism exerts on the liquid are distributed as a set of *N_k_* = 10 points over the body, *k* ∈ {±1, ±2*,…*, ±5}, with force positions determined by the contraction dynamics (1–2), which gives

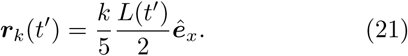

Next, the forces are estimated by the Stokes law,

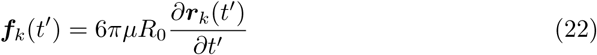

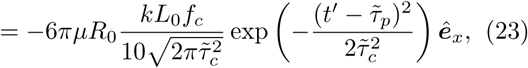

where the effective organism width *R*_0_ ≈ 100*µ*m. Finally, assembling Eqs. (20–23) provides a complete approximation for the contraction flow. The memory kernel convolved with the Gaussian force is readily integrated numerically. This solution is shown in Fig. S2, for high (1000x water) and low (water) viscosities, but with contraction time *τ_c_* constant.

#### C. Mixing

To analyse the mixing potential of the consecutive contraction and relaxation, we consider an ensemble of randomly distributed tracer particles in the organism vicinity, |***r****_p_*| *<* 5mm. The contraction flow is defined as in (20). The relaxation flow is given by the same equations, but with reversed contraction kinematics and much slower, with 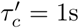. We integrate these numerically for a period of *t_f_* = 10s, using a fourth order Runge-Kutta scheme. The resulting tracer trajectories are shown in Fig. 2F of the main text. We then compute the final displacement after contraction and relaxation,

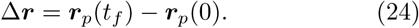

For the particles in 3% methyl cellulose, with kinematic viscosity *ν* = 50mm^2^*/*s, the hydrodynamics is dominated by viscosity, so the contraction and relaxation are reciprocal in time. In agreement with the scallop theorem [24], the forward trajectories are the same as the backward route, so the final displacement is negligible. For the particles in water however, with *ν* = 1mm^2^*/*s, this symmetry is broken and substantial mixing is observed, up to Δ***r*** ≈ 0.1*L*_0_.

### IV. RHEOSENSING

#### A. Experimental details

To measure their sensitivity to flows, organisms are tested in a novel microfluidic device that exerts hydrodynamic stresses on the cell body [Fig. S3]. The device is composed of two parallel disks (light blue) separated by a spacer (red), encased in an upright cylinder (gray) that holds liquid with organisms (dark blue). Organisms enter the flow chamber between the disks through four large inlets, and leave by the central outlet that is connected to a microfluidic pump. The chamber height is *H* = (700 ± 20)*µ*m, the inner radius *ρ*_1_ = 1mm and the outer radius *ρ*_2_ = 4cm. The disks are made of transparent acrylic that are imaged through with an inverted microscope (Nikon Te2000, 2X objective).

The exact solution of the flow pattern inside the chamber is known, the axisymmetric Jeffery-Hamel flow

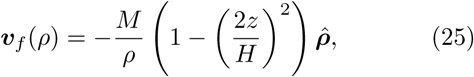

in cylindrical coordinates, with

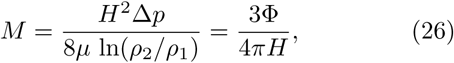

where Δ*p* is the pressure drop and Φ = (2.4 ± 0.1)cm^3^/s is the calibrated flow rate.

In this design, the flow increases with decreasing distance from the outlet, v*_f_* (*ρ*) ∝ −1*/ρ*, so while organisms are drawn towards the centre of the device they experience increasingly more hydrodynamic stress. The rate-of-strain tensor is defined in Cartesian components as

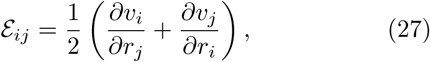

with *i, j* ∈{*x, y, z*} and the strain rate is defined as

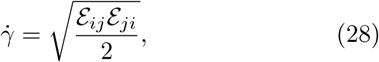

where the repeated indexes are summer over. Therefore, the strain rate in the middle of the flow chamber (25) is

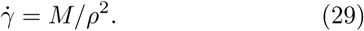

Thus, as an organism is drawn closer to the central outlet, the strain rate increases. When a critical value is reached, 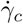, it contracts. This is observed around the critical radius, or the ‘event horizon’

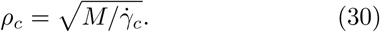

An additional advantage of this set-up is that elongated organisms naturally align with the radial direction, 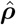, because they perform Jeffery-like trajectories with enhanced the time spent in the shear-extensional orientation [25, 26].

#### B. Analogy with tidal forces

Interestingly, one can make a direct analogy between the stretching of organisms in this suction flow device and ‘spaghettification’ near a black hole [27], a stretching induced by tidal forces due to non-homogeneous gravitational (cf. hydrodynamic) fields. Specifically, the Newtonian gravitational force on a body can be written as

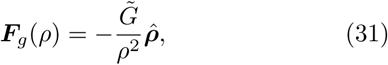

where 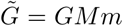 is the gravitational prefactor and *ρ* is the radial distance. A body of length *L* aligned along 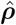 then experiences different forces at its head and tail ends, being

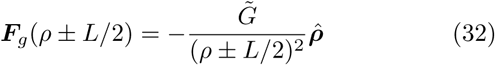

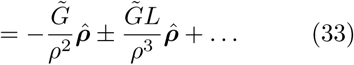

Here, the difference between the head and tail, the last term, gives the tidal force 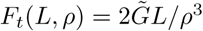.

Equivalently, the Jeffery-Hamel flow (25) is also nonhomogeneous, decaying with distance *ρ* as 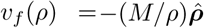, as evaluated at *z* = 0. Therefore, it also induces a difference in flow strength at the head and tail, giving the ‘hydrodynamic tidal force’

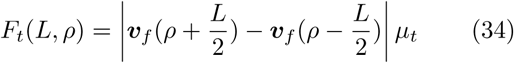

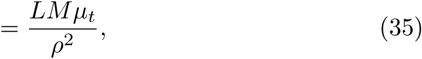

where *µ_t_* is a resistance factor, proportional to the fluid viscosity, that links the velocity with force. This expression can be compared directly to the strain rate (29). Also notice that it increases with cell length, *L*, which already suggests that longer cells can be better sensors. To understand this better, and how it depends on the organism shape, we study the distribution of tension that this tidal flow induces on the cell membrane, as discussed in the next section.

**FIG. S3.**
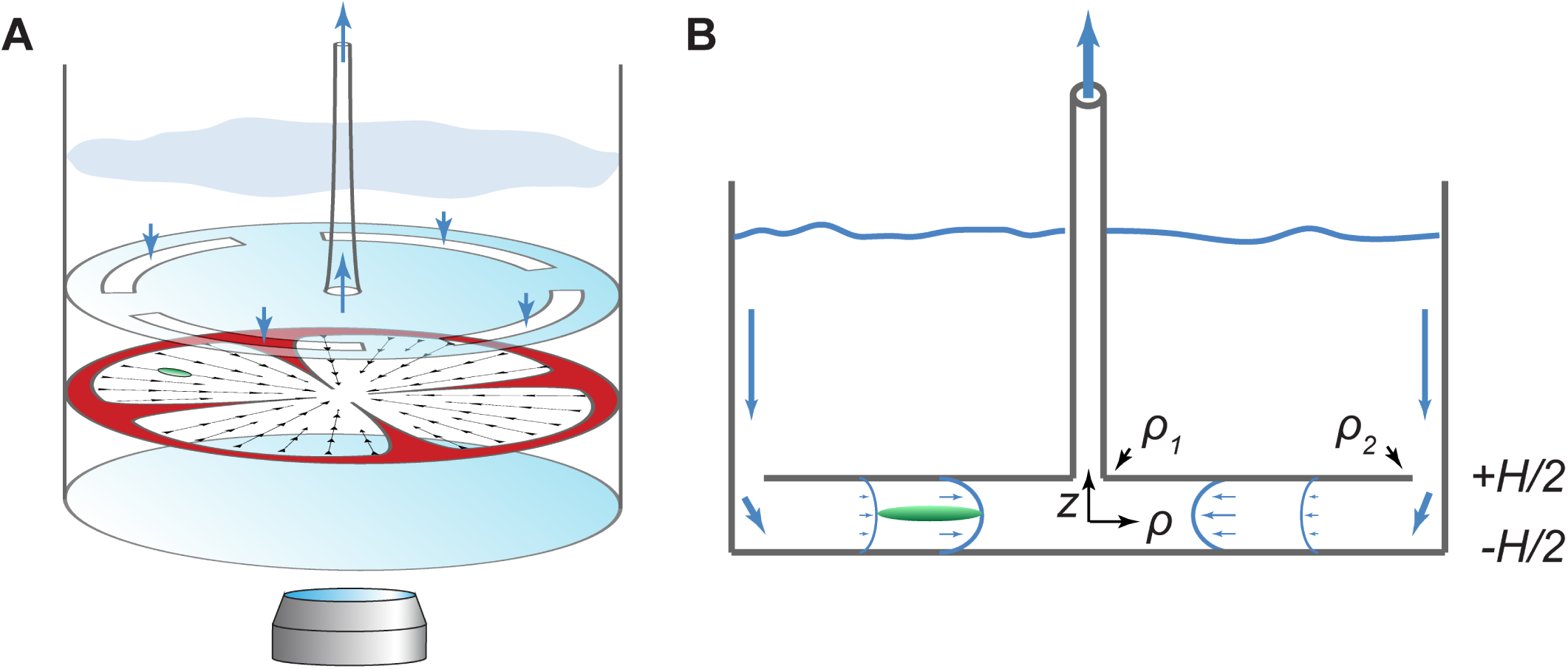
Rheosensing device (not to scale). Liquid with suspended organisms is held in a cylindrical container, with a narrow flow chamber in the bottom compartment. Liquid is drawn into the chamber on the sides and pumped out from a central tube. This creates an axisymmetric Jeffery-Hamel flow (25). A. 3D diagram of this experimental set-up. B. Side view, and schematic of flow profile. The organism (green) is stretched because the anterior flow is stronger than the posterior (28).

#### C. MRS calculations

Because the flow profile is known analytically in this device, it is possible to back-calculate the full distribution of forces that the liquid exerts on the membrane of the organism. We use the Method of Regularised Stokeslets (MRS) developed by Cortez et al. [28]. As the name implies, instead of the singular Oseen tensor (Eq. 7), a regularised tensor is introduced

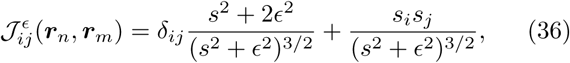

where ***s*** = ***r****_n_* − ***r****_m_*, *s* = |***s***| and ∊ is the regularisation parameter.

To account for the elongated *Spriostomum* shape, its surface is discretised with *N* points. We use the axisymmetric grid point distribution

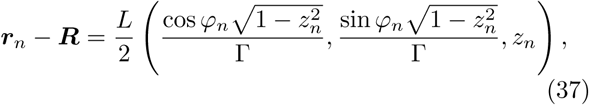

where we measure, for each individual organism, the centroid position just before contraction ***R***, the cell length *L* ~ 1mm, and the aspect ratio Γ ~ 10. We use *N_φ_* = 25 and *N_z_* = 100 so *N* = 2500, and

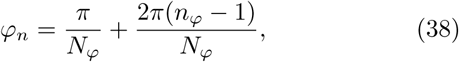

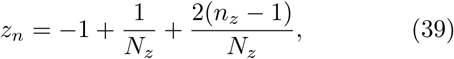

where the indexes *n_φ_, m_φ_* ∈ {1, 2*,…,N_φ_*} and *n_z_,m_z_* ∈ {1, 2*,…,N_z_*}. For this geometry, the regularisation parameter is set to 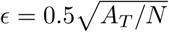, comparable to the distance between grid points, and defined in terms of the total surface area *A_T_* =2.8 10^−7^m^2^.

Now, the velocity of each point *n* on the organism can be decomposed in three components

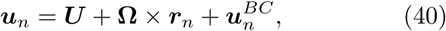

a rigid body translation ***U***, a rigid body rotation Ω and a surface deformation due to the applied flow, the boundary condition 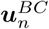. As in Eq. 8, these velocities are linked to the forces ***f****_m_* that the organism exerts on the liquid,

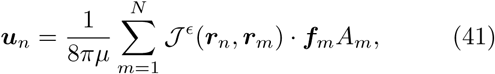

where *µ* is the fluid viscosity and *A_m_* are the quadrature weights,

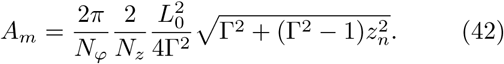

To find the forces that the liquid exerts on the membrane (−***f****_m_*), we must invert the linear system (41) subject to the constraints that the net force and torque must add up to zero in the absence of external contributions,

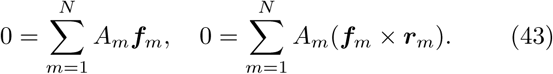

The boundary conditions are set by equating the flow inside the microfluidic device (25) to the surface velocity

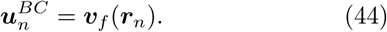

Therefore, we have 3*N* + 6 equations (41–44) and the same number of unknowns (***f****_n_*, ***U***, Ω).

This system is solved numerically for each organism at the moment just before contraction. Since we measure the positions ***r****_n_*, we compute the flows at each grid point ***v****_f_* (***r****_n_*) and then find the forces ***f****_n_* [black arrows in Fig. 3C]. Once the forces [N] are known, we compute the surface tension [N/m] along the long axis of the organism by dividing the summed forces over the surface cross section

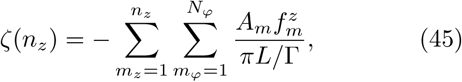

where Γ is the aspect ratio and *L/*Γ is the organism width. At the organism ends the membrane tension vanishes, ζ(0) = ζ(*N_z_*) = 0, due to the constraint (43). However, for a certain value *n_z_* halfway the tension peaks [colours in Fig. 3C]. Thus, we find the quantity of interest: the critical membrane tension, *T_c_* = max*_n_*_z_ ζ(*n_z_*), for each measured organism.

#### D. Ion channel theory

The membrane tension changes the probability that a mechanosensitive ion channel opens. To estimate this probability, we refer to the ‘two-state’ model by Phillips et al. [29], page 261 of the first edition. The channel can reside in the open (*σ* = 1) or closed (*σ* = 0) state. The energy of this system can be written as

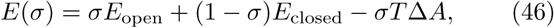

where the energies of the open and closed states are *E*_open_ and *E*_closed_, respectively, so Δ*E* = *E*_closed_ −*E*_open_ [J], the applied membrane tension is *T* [N/m] and the change in area upon gating is Δ*A* [m^2^]. The last term in Eq. 46 favours the open state, and reflects the work done to pull open the channel. The probability of finding the channel in a state with energy *E* is then given by the Boltzmann distribution, 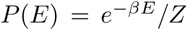, where 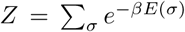 is the partition function and β =1*/k_B_T*. Therefore, the channel gating probability as a function of applied tension is

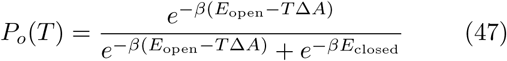

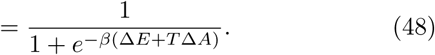

This function is plotted in Fig. 3D with Δ*E* = −5*k_B_T* and Δ*A* = (10nm)^2^. The critical membrane tension corresponds to the inflection point, *T_c_* = −Δ*E/*Δ*A* ≈ 0.2mN/m, which compares well with the measured value 〈*T_c_*〉≈ 0.17mN/m.

#### E. Strain-rate calculation for cell sizes

Instead of computing the membrane tension from the applied flow (§ IV C), conversely, it is also possible to compute the critical strain rate 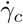 given the measured threshold tension 〈*T_c_*〉 = (0.17 ± 0.02)mN/m. This can be also done for different organism geometries. In particular, to answer the question what makes a better sensor, we are interested in 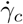 as a function of the cell size *L*.

We model this by using the Method of Regularised Stokeslets (MRS) once more. A virtual organism of length *L* is discretised (37) and placed in the flow (25) with virtual strength *M_v_*. The cell is oriented in the flow direction 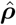 with its centroid at radius *ρ_c_*, so the virtual strain rate is 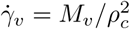. Exactly as before, we compute the forces ***f****_m_* and find the virtual maximum membrane tension *T_v_*. Then, we find the real critical flow rate *M_c_* required to reach the known critical tension *T_c_*, using the fact that the forces and the flow are linearly connected, 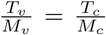, so we find 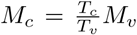. Hence, the critical strain rate required to trigger an organism of length *L* is

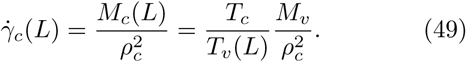

Note that the baseline flow *M_v_* and position *ρ_c_* are arbitrary, so we chose the measured experimental values. The resulting prediction is shown in Fig. 3E (black line).

### V. HYDRODYNAMIC TRIGGER WAVES

#### A. Clustering

We observe a robust clustering behaviour in *Spirostomum*, both in lab and in field conditions. After releasing organisms in a petri dish, the cells swim towards one another over typical time scales of a few minutes [Movie S7]. This accumulation is very consistent, even after manual fluid mixing.

The clustering mechanism is not well understood, but likely contributors are chemotaxis or biophysical pathways leading to motility-induced phase separation (MIPS) [30]. As organisms come close together, they could sense each other’s flows produced by swimming. If these small stimuli lead to ciliary reversal [Fig. S1B], then the mean swimming speed is reduced with increasing organism density v*_s_* ∝ 1*/n*, leading to dynamical clustering. Future research should look into this further.

**FIG. S4.**
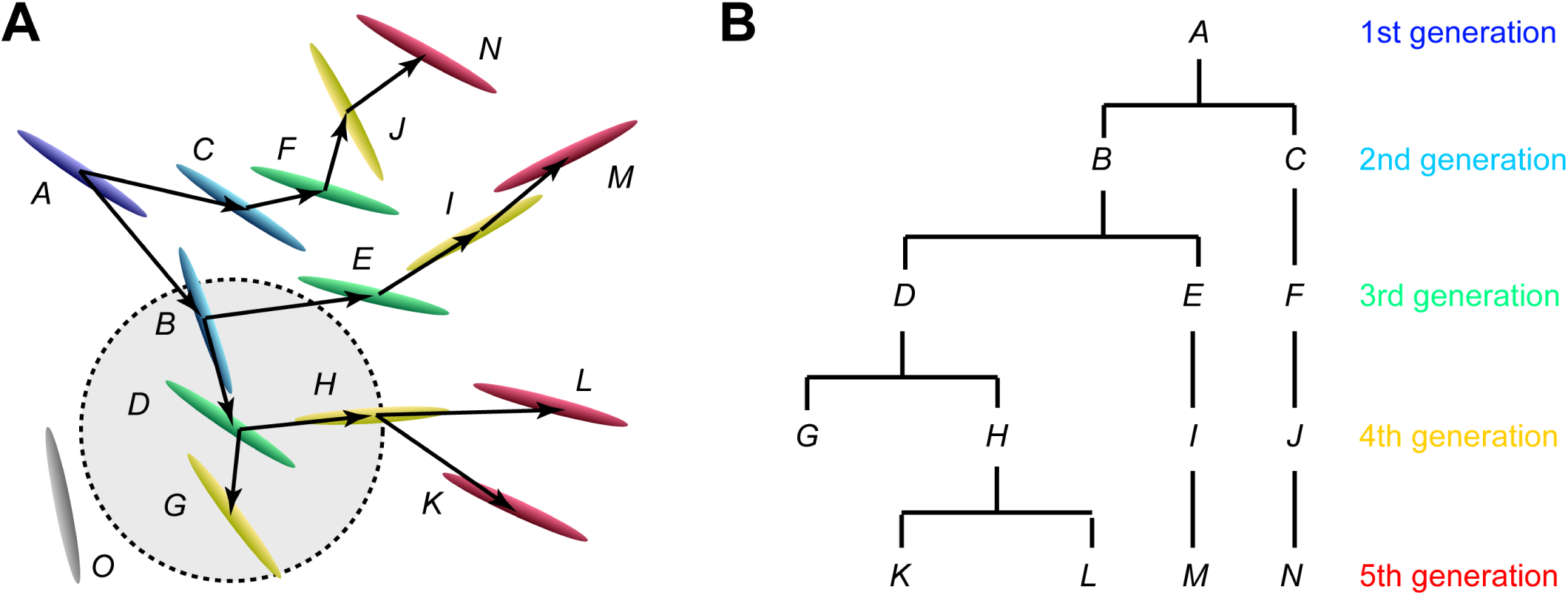
Collective hydrodynamic signalling. A. Example of configuration of organisms *A* − *O*. The first cell to contract is *A* (blue), and the last are *K* − *N* (red). One organism is not triggered, *O* (gray). The circle is used to compute the organism density around cell *D* (green). B. Connectivity graph of the same configuration.

#### B. Group contraction experiments

Collective dynamics were examined with high-speed (15002 fps) imaging of a dense (self-concentrated) organism suspension, gently transferred into a channel of parallel glass slides. The channel height, *H* = 200*µ*m, is not large compared to the cell width, *W* ~ 100*µ*m, to ensure the dynamics are primarily two-dimensional. Note that 3D group contractions are readily seen in petri dishes or culture flasks, but visualisation in 2D is more controlled. No electrical or other external stimuli were used here, but instead we waited until one organism would contract ‘spontaneously’, because of collision or flows generated by swimming. The resulting cascade then triggers a large group, ~ 100 cells in the field of view.

Recordings are analysed with a custom MATLAB script that (1) detects organism positions and orientations, (2) detects contraction events, and (3) deduces which organism triggered which, based on topography and timing, with statistical maximum likelihood.

From this information we construct a ‘connectivity graph’ that encodes each organism as a ‘vertex’ and each hydrodynamic signal as an ‘edge’ of the graph. Like a family tree, contractions are passed on from one generation to the next [Fig. S4]. At each signal event we extract the number density, *n* = *N/*(*πR*^2^), where *N* is the number of cells within a radius *R* = 1mm. We also find the time between contractions of each graph edge, 〈*τ_e_*〉 = (2.5±0.3)ms, which is indeed comparable with the electrical perception time measured in § II. Then, given the the centroid distance between sender and receiver, *ρ_e_*, we compute the trigger wave speed, 〈*v_w_*〉 = 〈*ρ_e_/τ_e_*〉. This trigger wave speed is plotted as a function of organism density [Fig. 4B].

#### C. Pair-wise interactions

Next to the above experiments at high organism densities (*n* ∈ [0.5, 5] cells/mm^2^), we also run low-density assays (globally ~ 0.1 cells/mm^2^). Here swimming cells encounter each other only occasionally in pairs, with a low probability that a third organism is nearby. We record long time lapses (8.93 fps, Leica DM4000 microscope, Hamamatsu camera) and detect pair-wise contractions with a custom MATLAB script. Thus, *N* = 79 interaction events were captured and analysed to extract the relative position (*ρ, θ*) and orientation (*ϕ*) between the transmitting and receiving organisms, (*A, B*), respectively [Fig. 4D,E, black stars].

We compare this to our theoretical model for flow generation and sensing. From the contraction flow (Eq. 20) we compute the strain rate 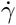 defined by Eq. 28 at the moment the flow is strongest, when 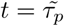 in Eq. 2. To keeps the model tractable, we consider the case without reflecting film surfaces (*m* = 0 only) and apply the Stokesian limit 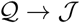. Both cell lengths are set to *L*_0_ = 1mm.

Figure 4C shows this strain rate 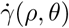 around the transmitting organism, *A*. Contours are shown for 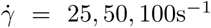 with green, orange and red lines, respectively, where the latter is the critical strain rate 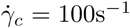. Then, we calculate the same strain rate at *N* points (37) along the body of the receiving organism B, for all configurations (*ρ, θ, ϕ*). In Fig. 4D this is plotted as a function of relative position, where the relative orientation and points on the body are maximised over;

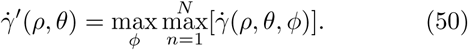

Similarly, the angular dependence is shown in Fig. 4E with fixed separation distance

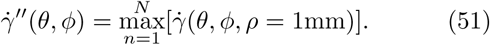

From this analysis we conclude that, indeed, the receiving organism *B* is most likely to contract when it crosses the red line, 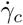, the ‘region of influence’.

#### D. Antenna theory

An analogy can be made here with electromagnetic antenna theory [31], where the distribution of output power in the (*θ*, φ) directions (in spherical coordinates) is given by the radiation intensity *U*(*θ*, φ) in units [Watt/steradian]. All real electromagnetic emitters and receivers are anisotropic, to satisfy the Helmholtz wave equation, enhanced by using parabolic dishes, which is often useful to transfer information in a specific direction. The directivity or directive gain is

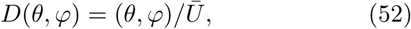

where 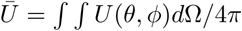 is the mean radiation intensity. The maximum directivity in any direction is then

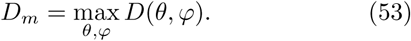

Thus, a value larger than 1 is considered gain. For example, a conventional half-wave dipole antenna has 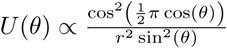, so the directive gain is *D_m_* =1.64 (2.15 dB).

*Spirostomum* also emits and receives hydrodynamic signals anisotropically, as shown in Fig. 4C. To approach this analytically we consider the Stokes dipole (10), which can be written in spherical coordinates as 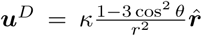 [32]. The corresponding strain rate (28) is then given by

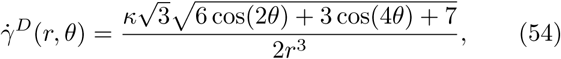

which like Fig. 4C also features two main lobes at *θ* =0*,π* and side lobes at *θ* = *π/*2. Hence, we obtain the dipole flow directive gain,

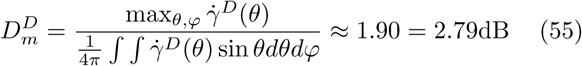

This quantifies that the hydrodynamic signals are transmitted ~ 2 times stronger in the direction of the long body axis. This dipole flow (10,54) is the most fundamental current generated by swimming micro-organisms [33]; e.g. also bacteria produce dipole flows [34].

#### E. Many-organism simulations

Building on this understanding, we continue to model multi-organism interactions by simulating large colonies of number density *n* ∈ [0, 5] mm^−2^. We consider an ensemble of *N* = *nπR*^2^ cells with length *L*_0_ = 1mm, distributed over a circle of radius *R_e_* = 20mm, with random orientations and positions. The first organism is set at the origin (0, 0) and starts contracting at *t* = 0. At each time step, for each non-triggered organism,

1. we sum the flows of the 6 nearest neighbours, if triggered at the previous time step. Further neighbour flows are not included to model the high fluid dissipation at high organism densities, a hydrodynamic screening effect [35, 36]. In other words, it would be unphysical for flows from further cells to travel through nearer cells.
2. From this flow we evaluate the maximum strain rate on points everywhere on the body, as for the pairwise-interactions.
3. If 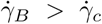, the receiving cells *B* are triggered. The duration between each time step is set by the perception time, which we model by the measured value Δ*t* = 〈*τ_e_*〉 =2.5ms.

We terminate the simulation if no more organisms are triggered, or if the signal reaches the colony edge, defined as radius *R_e_* − *L*_0_. If the latter happens, the ensemble reached percolation. The percolating path, the first connection from the origin to the edge, is shown as a coloured line in Figs. 4F-H. For each organism density *n* we repeat this simulation for *N_e_* = 100 ensembles, and compute the percolation probability *P* (*n*) as the fraction of percolating ensembles. We then also compute the mean wave speed 〈*v_w_*〉 = 〈*ρ_e_*〉*/*Δ*t*. Both quantities are plotted in Fig. 4B. The percolation threshold, where *P* (*n*) transitions from zero to unity, is found at 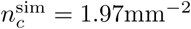.

#### F. Percolation theory

A theoretical estimate of the critical number density, *n_c_*, above which the hydrodynamic trigger waves can be sustained, is derived from percolation theory [37, 38]. This theory developed into the key framework for understanding conduction, signal transduction and long-ranged connectivity [39, 40], with many applications in biological systems [41], possibly including the dog alert in the Disney cartoon ‘101 Dalmatians’.

Because the organisms are distributed randomly in space, we consider continuum rather than lattice percolation. The classical continuum model is the formation of clusters by overlapping circles in 2D. The clusters get larger as the number of circles increases, or specifically as the fractional area that they cover *p* ∈ [0, 1] grows. Then, at a critical value *p_c_* ≈ 0.676 a ‘spanning cluster’ emerges that is as large as the system size. For this problem, an exact series expansion solution was found [42].

Because we are interested in the non-circular structure of the *Spirostomum* signals, we continue by considering ellipses. An analytical approximation for overlapping ellipses was published by Yi et al. [43], and an interpolation formula was found by Xia et al. [44], giving the critical area fraction

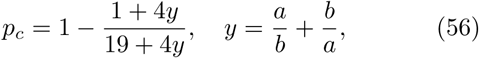

in terms of the major and minor semiaxes, *a>b*. Given the area of an ellipse, *A* = *abπ*, the relationship [44] between the non-dimensional area fraction *p* and the dimensional number density *n* is

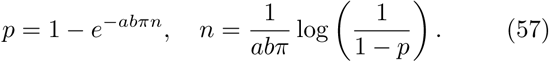

Instead of the organisms themselves overlapping, we consider the *region of influence* of the transmitting organisms’ flows (see § V C) crossing the bodies of the receiving organisms. Therefore, we use the mapping

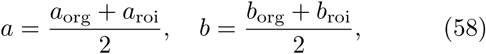

where *a*_org_ = 10*b*_org_ =0.5mm and we obtain (*a*_roi_, *b*_roi_) ≈ (0.863, 0.302)mm from fitting an ellipse to the region of influence [Fig. 4C, dashed red line]. Inserting (58) into (56, 57) then gives an estimate for critical organism density, 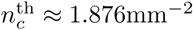.

A further analytical estimation without any fitting parameters can be obtained by approximating the region of influence using the dipole approximation (54) for the strain rate. Solving 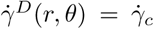 for the radius *r* at *θ* = (0, *π/*2) gives

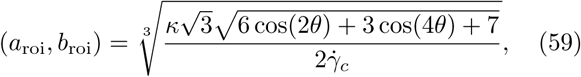

where the dipole moment *κ* = *f*_max_*L*_0_*/*8*πµ* ≈ 5.98mm^3^*/*s and 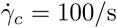. Inserting these values gives (*a*_roi_, *b*_roi_) ≈ (0.592, 0.470)mm, leading to the critical organism density, 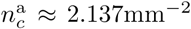. This expression can be applied directly to other organisms for which the flow (or dipole moment) and critical strain rate are known.

The presented percolation theory can be extended straightforwardly to more natural 3D organism colonies using overlapping ellipsoids of revolution [45]. On a more fundamental level, we expect the hydrodynamic signals to be part of the *directed percolation* universality class [46–48]. Here the directionality can be interpreted as a temporal degree of freedom [40], like following ‘one-way streets’ in a maze. Analogously, the organism signals also have uni-directional paths for certain relative configurations (*ρ, θ, ϕ*), where a cell can trigger a neighbour but that neighbour cannot trigger the former. An example is 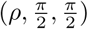 versus 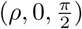 [Fig. 4E]. Therefore, the physical properties of the phase transition (the critical exponents) might be different from the overlapping shapes, a member of the ‘ordinary percolation’ universality class. This could be an exciting new avenue for future research in hydrodynamic communication.

### VI. LIST OF SUPPLEMENTARY MOVIES

- Movie S1.mov – Close-up of metachronal waves, antisymplectic on top. Images are recorded at 40x magnification and 500fps (playback 10x slowed down). The organism swims between glass slides separated 100 microns. Added are *D* =0.75 micron tracer particles to visualise the local flows generated by the swimming strokes.
- Movie S2.mov – Montage of global flows generated by swimming. Images are recorded at 4x magnification and 500fps (playback 10x slowed down). The organism swims in a glass channel of height 100 microns, filled with *D* =0.75 micron tracer particles. The FlowTrace algorithm [14] was used to convolve a superposition of 100 frames for each image.
- Movie S3.mov – High-speed recording of body dynamics during contraction. Images are recorded at 20x magnification and 2,800fps (playback 50x slowed down). The contraction occurred ‘spontaneously’, so was not triggered electrically.
- Movie S4.mov – High-speed recording of inertial flows generated during a contraction. Images are recorded at 10x magnification and 10,000fps (playback 200x slowed down). The organism swims in an air-liquid-air film of 500 microns height, and was triggered by an electric pulse (|***E***| =1.5kV/m, *τ_E_* = 100ms) in the middle of the film. The Flow-Trace algorithm [14] was used to show streamlines by superimposing 200 frames for each image. Note the spreading of the 4 vortex centers into the liquid over time, cf. Movie S5.
- Movie S5.mov – Linearised Navier-Stokes solution of inertial flows generated during a contraction. The flow solution (Eq. 20) is evaluated in a 2×2mm domain during a 100ms period. The method of images was used to satisfy the boundary conditions of an air-liquid-air film of 500 microns height. Yellow arrows are streamlines and background colours denote the flow magnitude. Note the spreading of 4 vortices into the liquid over time, cf. Movie S4.
- Movie S6.mov – Rheosensitivity measured in a microfluidic device. Images are recorded at 1.5x magnification and 1,000fps (playback 50x slowed down). As organisms are drawn towards the central flow outlet (red), the strain rate increases with decreasing radial position, 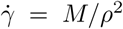, where *M* is the flow rate. When the organism reaches the threshold (orange, 〈*ρ_c_*〉 =3.22mm) its mechanosensitive channels open and the cell contracts. This high-throughput experiment is repeated for *N>* 100 cells to compute the average critical strain rate.
- Movie S7.mov – Spontaneous clustering behaviour. Images are recorded at 1x magnification and 4fps (playback 10x speed up). The organisms swim towards one another after manual fluid mixing. A double group contraction is observed at cell high densities, at time point 5:30.
- Movie S8.mov – Collective hydrodynamic signaling. Images are recorded at 4x magnification and 15,000fps (playback 150x slowed down). Contracting cells generate a flow that trigger other organisms. Superimposed on the experimental video is the evolution of the connectivity graph, showing which organism triggered which, where colours denote the contraction time.
- Movie S9.mov – Percolation simulations of collective hydrodynamic signaling. A large number *N* = *nπR*^2^ ~ 15, 000 of organisms (black lines) are distributed randomly over a circular cluster of radius *R* = 50mm. One central organism is triggered, and subsequent contraction dynamics are simulated over time (denoted by colours) until no more organisms move. (a) Just below the critical point, at density *n* = 1.75 [cells/mm^2^], the trigger wave decays very rapidly. (b) Near the critical point, at *n* = 2, many branches of the connectivity graph die out but one path reaches the cluster edge. (c) Just above the critical point, at *n* = 2.25, almost all organisms are triggered, readily establishing percolation. In summary, a 25% difference in density has a much larger effect on communication.
- Movie_S10.mov - Simplified demonstration of wave speed dependence on organism density. Cells are arranged along the x axis with parallel (a) and perpendicular (b) orientations, separated a distance just small enough for signal transduction. At lower cell densities (a) the wave moves faster than higher densities (b), but at the cost of losing percolation by small orientation fluctuations. In this simplified case the cells can only trigger the nearest neighbours. In more complex configurations the hydrodynamic screening at high cell densities has a similar effect

